# Fast and effective protein model refinement by deep graph neural networks

**DOI:** 10.1101/2020.12.10.419994

**Authors:** Xiaoyang Jing, Jinbo Xu

**Affiliations:** Toyota Technological Institute at Chicago, Chicago, IL 60637, USA

## Abstract

Protein structure prediction has been greatly improved, but there are still a good portion of predicted models that do not have very high quality. Protein model refinement is one of the methods that may further improve model quality. Nevertheless, it is very challenging to refine a protein model towards better quality. Currently the most successful refinement methods rely on extensive conformation sampling and thus, take hours or days to refine even a single protein model. Here we propose a fast and effective method that may refine protein models with very limited conformation sampling. Our method applies GNN (graph neural networks) to predict refined inter-atom distance probability distribution from an initial model and then rebuilds the model using the predicted distance as restraints. On the CASP13 refinement targets our method may refine models with comparable quality as the two leading human groups (Feig and Baker) and greatly outperforms the others. On the CASP14 refinement targets our method is only second to Feig’s method, comparable to Baker’s method and much better than the others (who worsened instead of improved model quality). Our method achieves this result by generating only 5 refined models for an initial model, which can be done in ∼15 minutes. Our study also shows that GNN performs much better than convolutional residual neural networks for protein model refinement when conformation sampling is limited.

**Availability:** The code will be released once the manuscript is published and available at http://raptorx.uchicago.edu

## Introduction

High-accuracy protein structure prediction can facilitate the understanding of biological processes at the molecular level. In the past few years, protein structure prediction has been greatly improved, mainly due to the introduction of deep convolutional residual networks (ResNet)^1–4^ and lately transformer-like networks implemented in AlphaFold2. However, a good percentage of predicted protein structural models still deviate from their native structures, which limits their value in downstream applications. To further improve model quality, much effort has been devoted into developing model refinement methods^5–7^. The main goal is to refine an initial model towards its native structure and then, to generate new models of higher quality. This is a very challenging task since the space of worse models is much larger than that of better models. Many refined models submitted by CASP participants have worse quality than their starting models^5^.

A typical model refinement method employs side-chain repacking, energy minimization and constrained structure sampling^8–11^. Since the energy function is usually challenging to optimize, model quality may not be improved without large-scale conformational sampling. Currently, the most successful refinement methods use large-scale conformational sampling either through molecular dynamics (MD) simulations^6^ or fragment assembly^7,12^. For example, Feig group employs iterative MD simulation with flat-bottom harmonic restraints to sample conformations. A subset of sampled models are selected using Rosetta scoring function and averaged to build the final refined model. Baker group^7^ uses local error estimation to guide conformational sampling by fragment assembly, and iteratively refines the models by recombining secondary structure segments and replacing torsional angles. The lowest-energy model in the last iteration is identified using Rosetta scoring function and then averaged with its conformational neighbours to build the final refined model. GalaxyRefine2^12^ developed by Seok group employs multiple conformation search strategies. Model error estimation can be used to constrain the sampling space and prevent degradation of the stable structure regions. DeepAccNet^13^ uses both 3D and 2D convolution networks to estimate residue-wise accuracy and inter-residue distance error, which are then converted into Rosetta restraints to guide conformational sampling. Although performing well on some proteins, these methods rely on extensive conformational sampling and thus, a lot of computing resources for even a single protein model ^13,14^.

In this work we propose a new model refinement method GNNRefine that may quickly improve model quality without extensive conformation sampling. GNNRefine represents an initial protein model as a graph and then employs graph neural networks (GNN) to refine it. GNN has been used to predict protein model quality^15,16^, but not to refine protein models. GNNRefine iteratively updates the node (residue-wise) and edge (residue-pair) features in the graph by multiple message-passing layers to capture the global structural information, from which it predicts inter-atom distance probability distribution. The predicted distance probability is converted into distance potential, which is then fed into PyRosetta^17^ FastRelax^18^ to produce refined models without extensive conformational sampling. Our experimental results show that on average GNNRefine may improve model quality, significantly outperforms those methods without using large-scale conformational sampling, and is slightly worse than Feig’s leading method that uses large-scale conformational sampling. Another advantage is that GNNRefine produces fewer degraded models than other methods.

## Results

### Overview of the method

Figure 1A shows the flowchart of our method GNNRefine, which mainly includes three steps: 1) represent the initial model as a graph and extract atom, residue, and geometric features from the initial model, 2) predict refined distance for each edge in the graph using graph neural network (GNN), and 3) convert the predicted distance probability into distance potential and feed it into PyRosetta^17^ FastRelax^18^ to produce refined models by side-chain packing and energy minimization. Meanwhile, the GNN-based distance prediction is the key factor affecting the refined model quality. As shown in Figure 1B, GNNRefine mainly consists of three modules: an atom embedding layer, multiple message passing layers, and an output layer. The atom embedding layer is used to learn atom-level structure information of one residue and the resultant atom embedding is concatenated with other residue features to form the final feature of a residue. The protein graph is built on the residue feature (node) and bond or contact feature (edge) between residue pairs (detailed in the Methods section). By going through multiple message passing layers, the node and edge features are iteratively updated to capture global structural information. Finally, a linear layer and a softmax function are used to predict distance probability distribution from the edge feature.

**Fig. 1.**
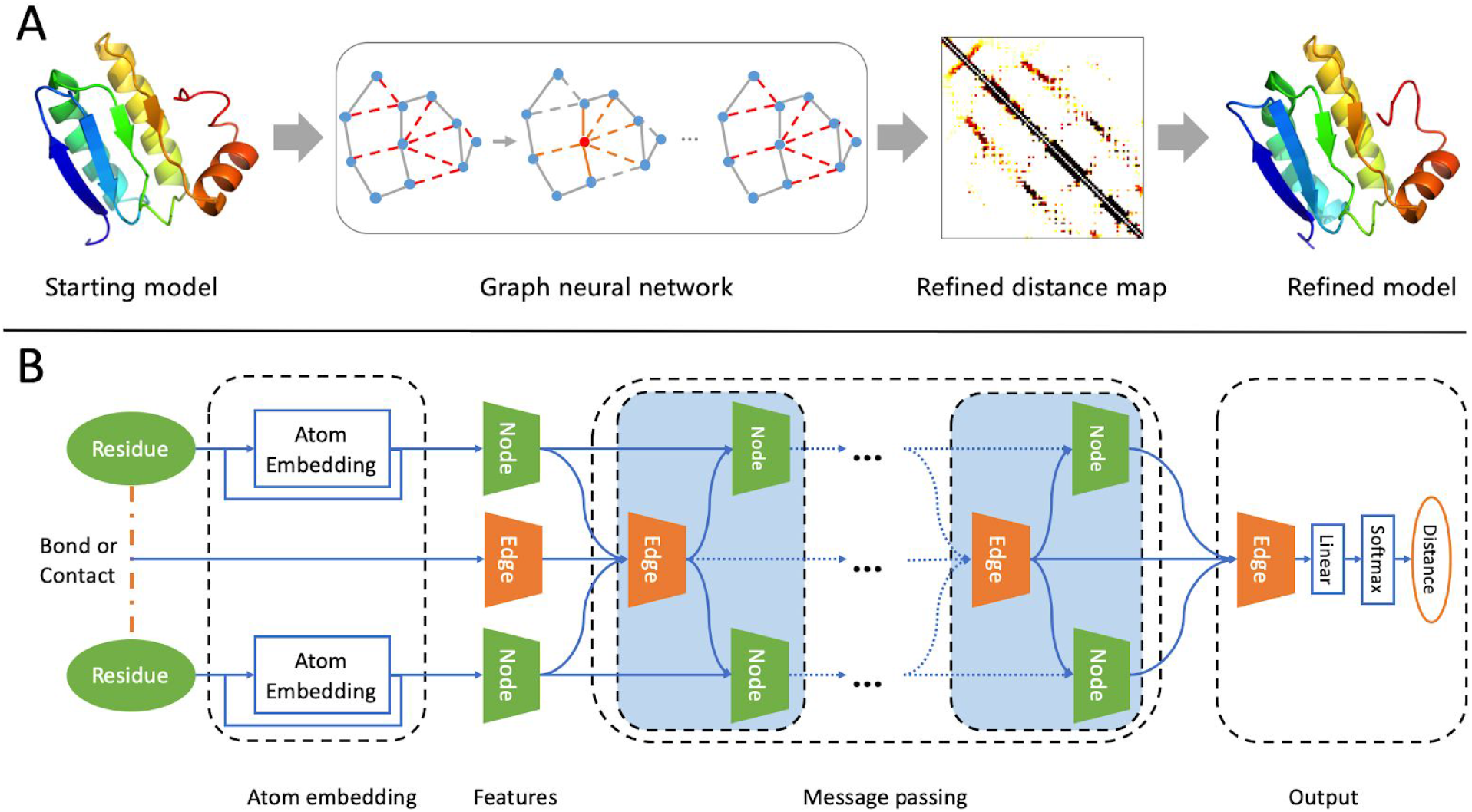
The GNNRefine method for protein model refinement. A. the flowchart of GNNRefine including feature extraction, refined distance prediction by GNN, and refined model building by FastRelax; B. the network architecture of GNNRefine.

The predicted distance probability is converted into distance potential which is then fed into PyRosetta FastRelax to build the refined model. Since we use a much smaller number of distance restraints (i.e., only consider those residue pairs with distance no more than 10Å in the starting model) and directly refine the model without a lot of samplings, our method runs very fast. Tested on the CASP13 dataset, our method needs on average only 15 minutes to refine a single protein model when 10 CPUs are used to run FastRelax (to generate 50 refined models). In contrast, Baker’s DeepAccNet needs more than 10 hours on 50 CPUs to refine a single model with 120 residues.

### Evaluation metrics

We evaluate quality improvement of the refined models over their starting models in terms of GDT-HA, GDT-TS and lDDT. We also use “Degradation” to count how many refined models have quality worse than their initial models by a given threshold (0, −1 and −2). Meanwhile, 0 denotes that a refined model has worse GDT-HA than its starting model; −1 and −2 denote that a refined model’s GDT-HA is worse than its starting model by at least 1 and 2 units, respectively.

### Performance on the CASP13 refinement targets

We compare our method with two leading human groups in the CASP13 refinement category^5^ (FEIGLAB and BAKER) and 5 server groups Seok-server, Bhattacharya-Server, YASARA, MUFold_server and 3DCNN. Their refined models are available at the CASP official website. A human group has up to three weeks to refine one model while a server group has at most three days. A human group may make use of any extra information. For example, FEIGLAB selected refined models manually and BAKER group chose their sampling strategy based upon the model quality provided by the CASP organizers. Here we evaluate the quality of the first submitted models, as shown in Table 1. Fig. 2 shows the box plot of the ΔGDT-HA distribution. Even if generating only 5 refined models for each initial model, our GNNRefine has comparable performance as the two human groups and outperforms all the 5 servers in terms of quality improvement. Seok-server is the only server that obtained positive improvement on the three metrics. Bhattacharya-Server and YASARA improved lDDT slightly, but degraded GDT-HA and GDT-TS. MUFold_server and 3DCNN degraded all the three metrics. Further, our method generates only 4 refined models with slightly worse quality than their initial models, but all the other methods including the two human groups degraded many models. That is, it is very safe to use our method to refine models. Fig. 2 shows that the two human groups have a larger variance than our method possibly because of extensive conformational sampling. That is, they may be able to refine some models very well, but also likely to degrade some models a lot. In contrast, our method has a smaller variance since we do not use extensive conformational sampling.

**Table 1.**
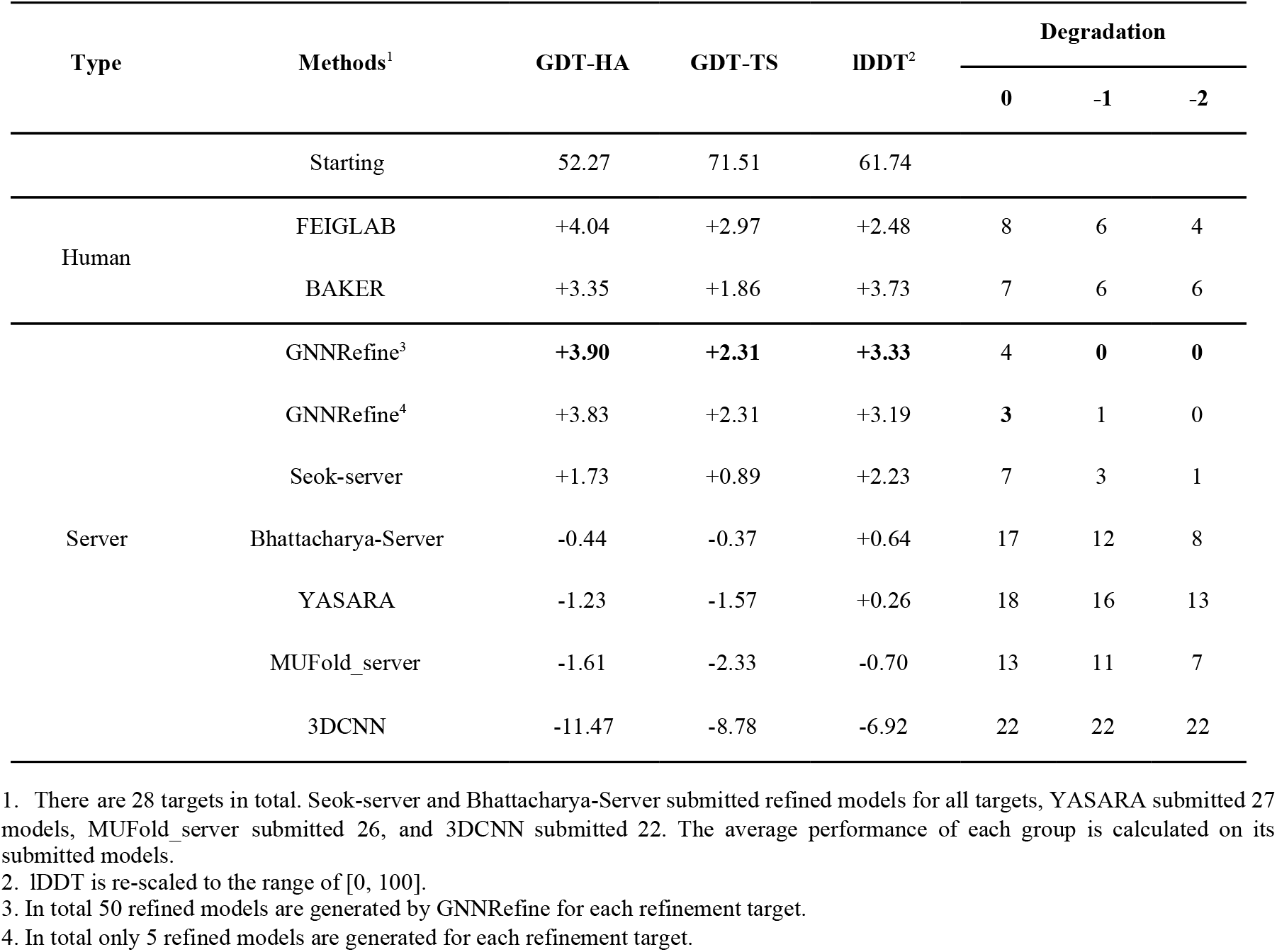
Performance on the CASP13 refinement targets

**Fig. 2.**
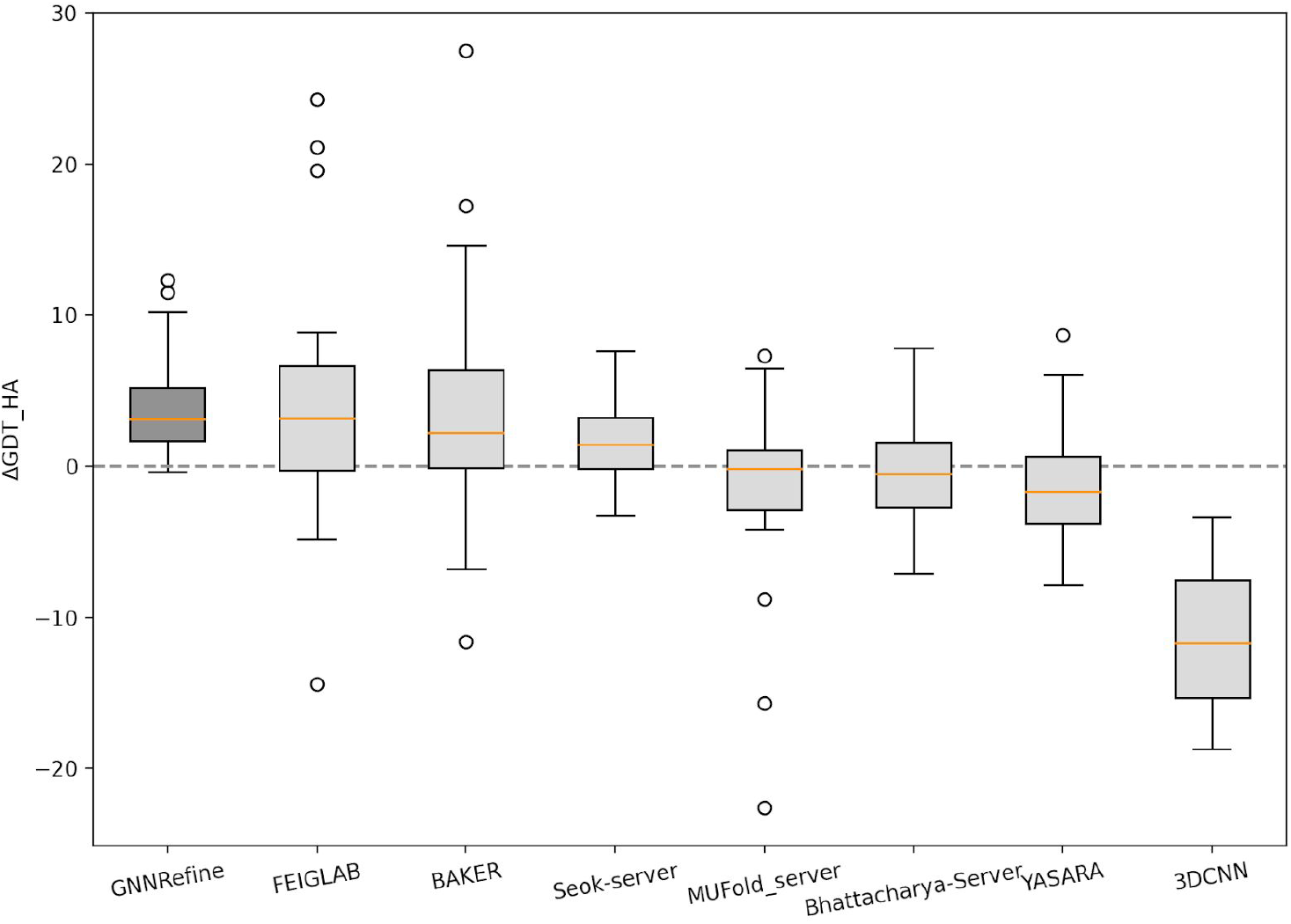
Box plot of the distribution of ΔGDT-HA values on the CASP13 refinement targets

### Performance on the CASP14 refinement targets

We test our method on the 37 CASP14 refinement targets and compare it with two human groups FEIG and BAKER and 4 server groups FEIG-S, Seok-server, Bhattacharya-Server and MUFold_server. It should be noted that we did not finish this work before CASP14, so our method was not blindly tested in CASP14. FEIG-S is a server group, but it is not fully automated for some targets as mentioned in the CASP14 abstract^19^. Table 2 summarizes the performance and Fig. 3 shows the box plot of the ΔGDT-HA distribution. The CASP14 models are much harder to refine than CASP13 models. All the server groups except FEIG-S degraded the model quality in terms of GDT-HA, GDT-TS and lDDT, and all including the two human groups degraded the quality of more than 10 models. This is because some initial models are well-refined, especially the 14 AlphaFold2 models. If excluding the AlphaFold2 models, the model quality improvement is comparable to CASP13 (Supplementary Table S1). Overall, on the CASP14 refinement targets, our method performs slightly worse than Feig’s methods, comparably to Baker’s method and better than the others. Our method degraded the least number of models. It is worth mentioning that even generating only 5 refined models for an initial model, our method does not lose refinement accuracy.

**Table 2.**
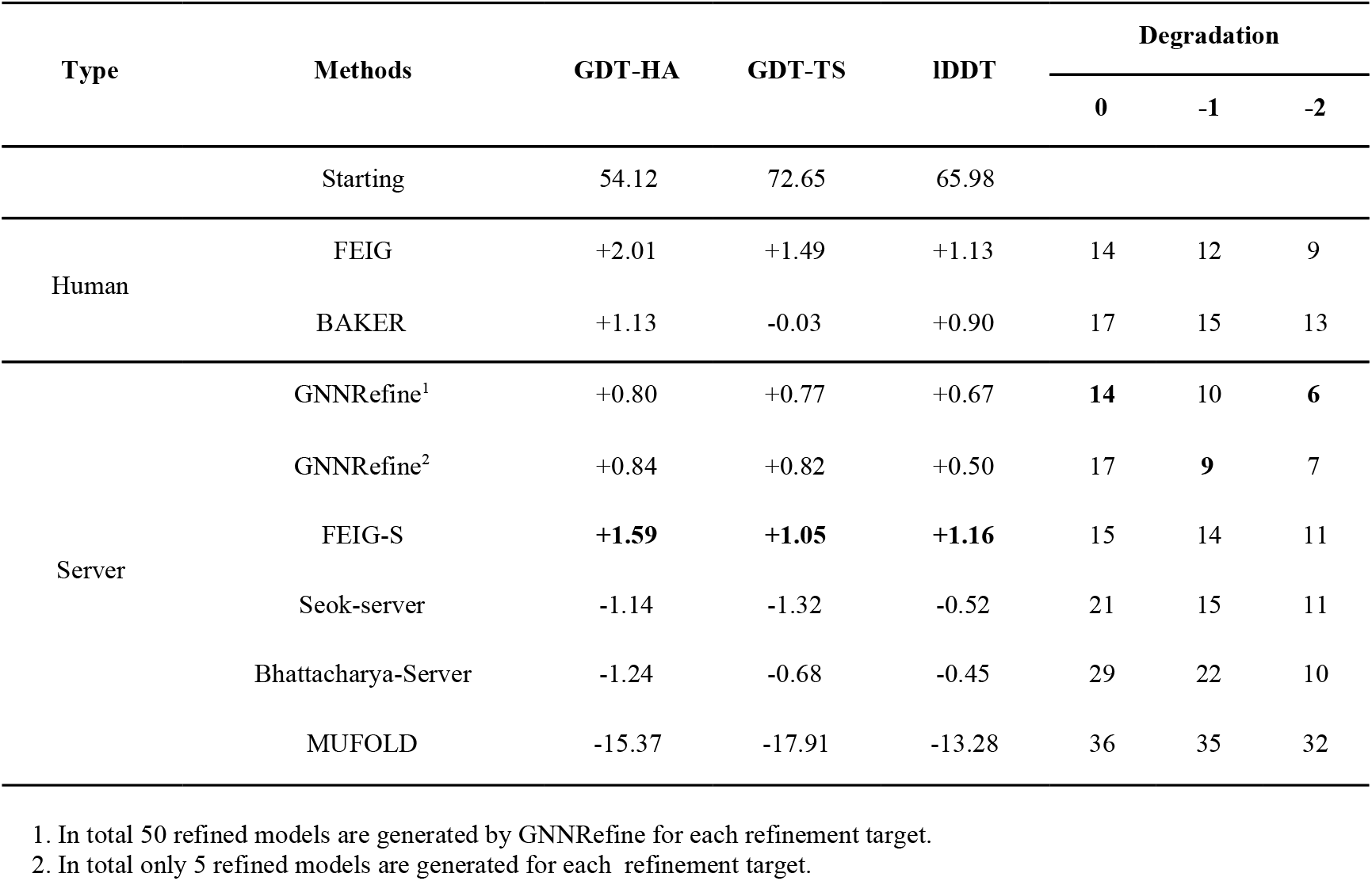
Performance on all CASP14 refinement targets

**Fig. 3.**
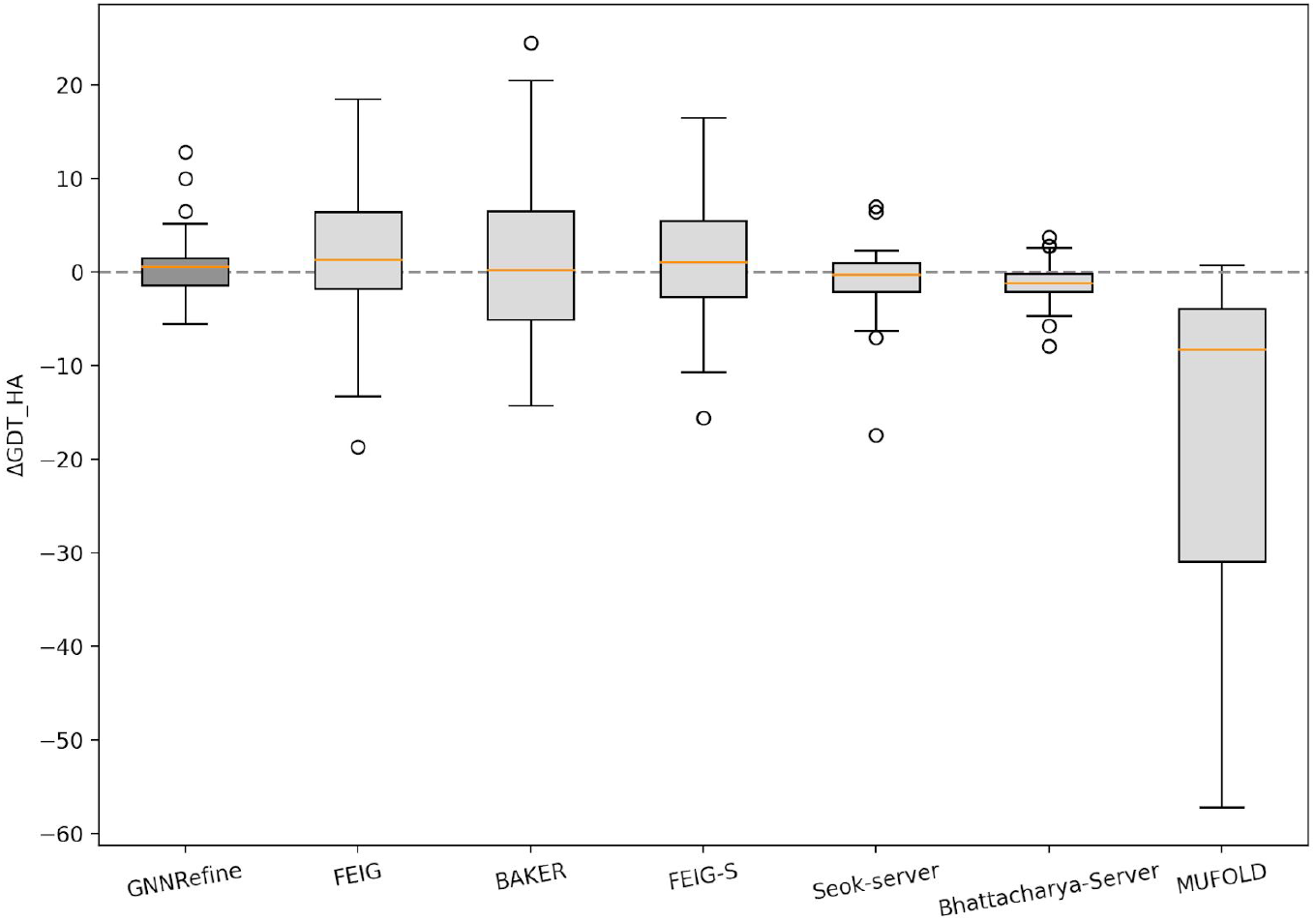
Box plot of the distribution of ΔGDT-HA values on the CASP14 refinement targets

GNNRefine has successfully refined five CASP targets (3 CASP13 targets and 2 CASP14 targets) with ΔGDT-HA ≥10. Fig. 4 shows 4 of them with publicly available experimental structures and indicates that our method can refine the starting model at different secondary structure regions (helix, sheet and coil).

**Fig. 4.**
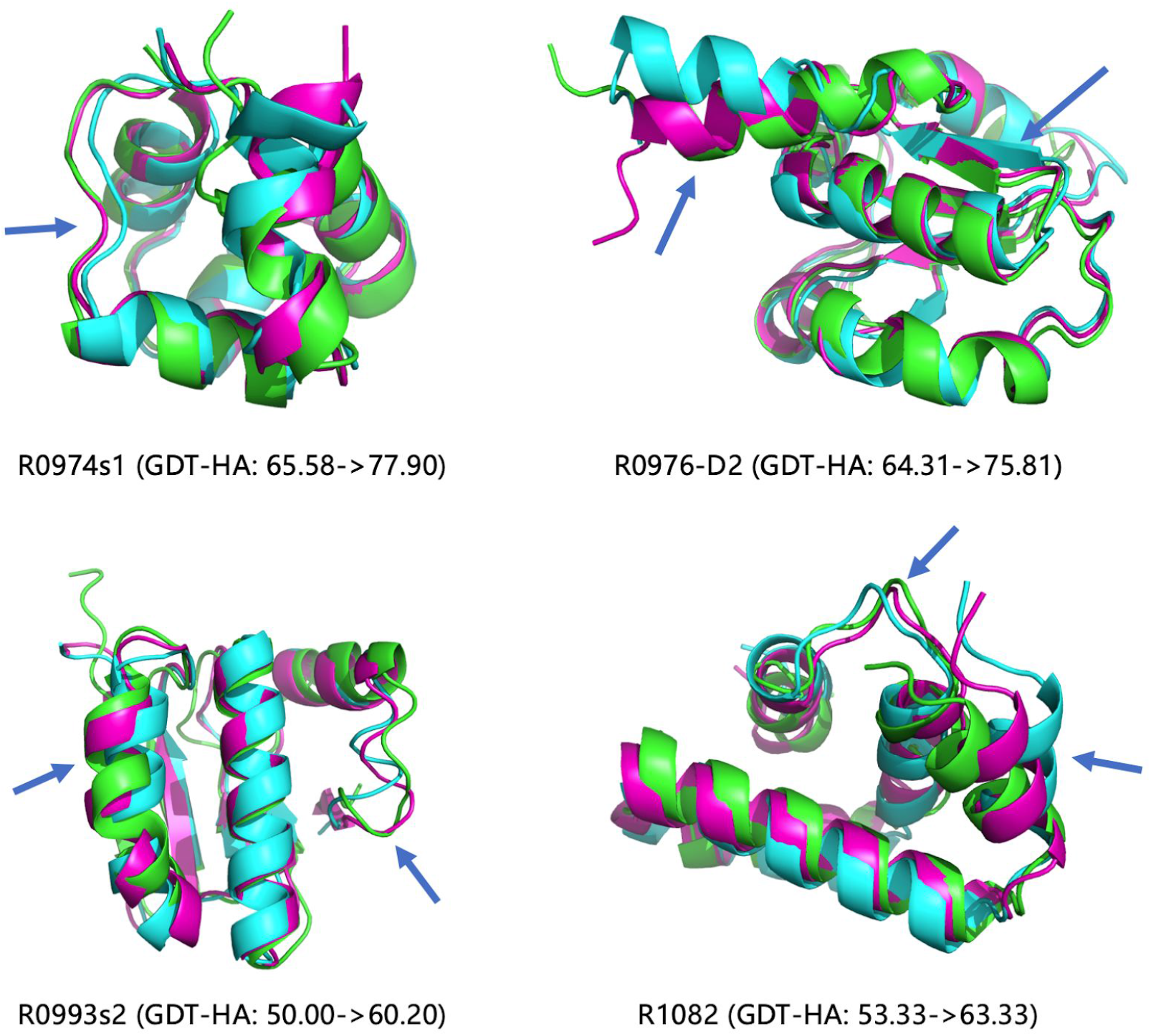
Successful refinement examples by GNNRefine for targets R0974s1, R0976-D2 and R0993s2 from CASP13, and R1082 from CASP14. Native structures, starting models, and refined models are shown in green, cyan, and magenta, respectively. Incorrect regions in the starting models that were significantly refined are indicated with blue arrows.

### GNNRefine outperforms existing standalone software

Here we compare our method with some publicly available software such as GalaxyRefine^9^ and ModRefiner^8^. We run GalaxyRefine locally by its default configuration. ModRefiner has a configurable parameter strength in [0, 100] to control the strength of restraints extracted from the starting model, with strength 0 meaning no restraints at all while strength 100 indicating very tight restraints by the starting model. We run ModRefiner with three different strength values: 0, 50 and 100. As a control, we also run PyRosetta FastRelax without using the distance restraints predicted by GNNRefine. Table 3 shows the performance and running time on the CASP13 targets, our method outperforms the other methods by all metrics.

**Table 3.** Performance of standalone software on the CASP13 refinement targets

| Methods | GDT-HA | GDT-TS | IDDT | Degradation |  |  | Running Time<sup>1</sup> |
| --- | --- | --- | --- | --- | --- | --- | --- |
|  |  |  |  | 0 | -1 | -2 |  |
| Starting | 52.27 | 71.51 | 61.74 |  |  |  |  |
| GNNRefine<sup>2</sup> | **+3.90** | **+2.31** | **+3.33** | 4 | **0** | **0** | ~2.5 (~0.25<sup>3</sup>) |
| GNNRefine<sup>4</sup> | +3.83 | +2.31 | +3.19 | **3** | 1 | 0 | ~0.25 |
| GalaxyRefine | +0.21 | +0.05 | +1.23 | 15 | 8 | 4 | ~2.42 |
| ModRefiner-100 | +0.16 | +0.05 | +0.73 | 13 | 3 | 0 | ~0.38 |
| ModRefiner-50 | -0.05 | +0.04 | +0.78 | 11 | 6 | 0 | ~0.47 |
| ModRefiner-0 | -0.70 | -0.58 | +0.99 | 17 | 12 | 2 | ~0.58 |
| FastRelax | -2.00 | -1.96 | +0.17 | 18 | 14 | 13 | ~0.05 |

1. The average running time (CPU hour) needed to refine a single protein model. 2. In total 50 refined models are generated by GNNRefine for each refinement target. 3. The running time is 2.5 hours on a single CPU and reduced to 0.25 hours when running in parallel on 10 CPUs. 4. In total only 5 refined models are generated for each refinement target.

### Performance on the CAMEO dataset and CASP13 FM dataset

We further evaluate the performance of our method on two large datasets. The first consists of 208 starting models for the CAMEO targets and the second consists of 4193 decoy models built by our in-house template-free modeling method for the CASP13 FM targets. See Supplementary Table S2 and S4 for the performance on the CAMEO targets and CASP13 FM targets. We also analyze the correlation between the quality of the starting model and the improvement by GNNRefine on the CAMEO targets, as shown in Supplementary Figure S1 and Table S3. The results show that our method performs better on high quality models, which means that our method has great potential for high-accuracy model refinement.

### GNNRefine improves distance prediction

To understand why GNNRefine works without extensive conformational sampling, we evaluate the distance predicted by GNNRefine in terms of top L contact precision and lDDT. For each residue pair, the predicted probabilities of distance bellow 8Å are summed up as predicted contact probability. We select top L contacts in the starting model by their respective C_β_-C_β_ Euclidean distance ascendingly. To calculate the lDDT of the distance predicted by GNNRefine, for each residue pair we use the middle point of the bin with the highest predicted probability as its real-valued distance prediction. We only consider the C_β_-C_β_ pairs with predicted distances less than 20Å. Table 4 shows that the distance predicted by GNNRefine is better than the starting model in terms of both contact precision and lDDT.

**Table 4.**
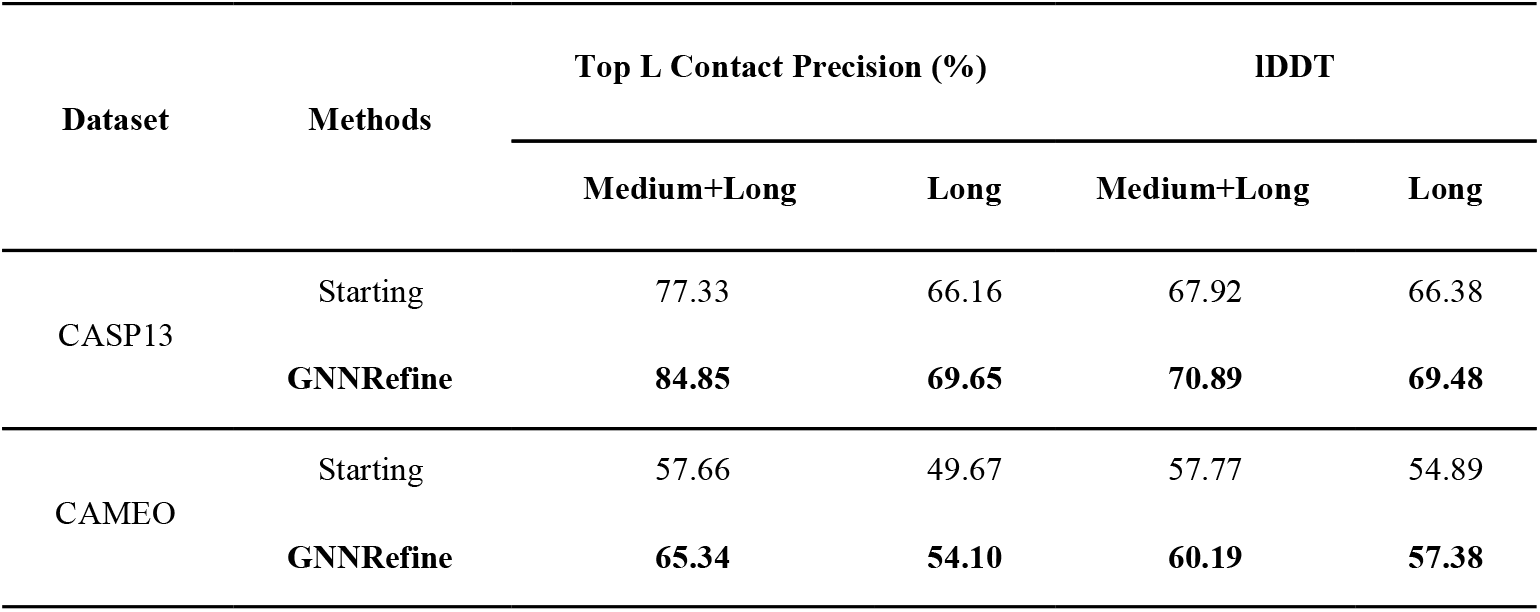
Comparison between predicted distances and distances in the starting model

### GNN outperforms ResNet for model refinement

The convolutional residual neural network (ResNet) is widely used for protein contact and distance prediction. Baker group developed a ResNet-based method DeepAccNet for model refinement. To test the performance of DeepAccNet with limited conformational sampling, we feed the distance potential generated by DeepAccNet into PyRosetta FastRelax to build refined models, using exactly the same method as GNNRefine. We have also developed an in-house ResNet model (of 41 2D convolutional layers) to predict distance from initial models and test if the distance predicted by it can be used to refine models or not. To compare the three methods fairly, we use only one deep GNNRefine model to do refinement in this experiment. For each method, we generate 10 refined models from each starting model and select the lowest-energy model as the final refined model. Table 5 shows that our GNN method greatly outperforms our in-house ResNet method, which in turn is better than DeepAccNet. That is, DeepAccNet is not able to refine models when extensive conformation sampling is not used, but our GNN method works. The difference between our in-house ResNet and DeepAccNet lies in that our ResNet directly predicts distance distribution while DeepAccNet predicts the distribution of distance error. Table S5 shows that our GNN method indeed can predict distance with better accuracy than ResNet.

**Table 5.**
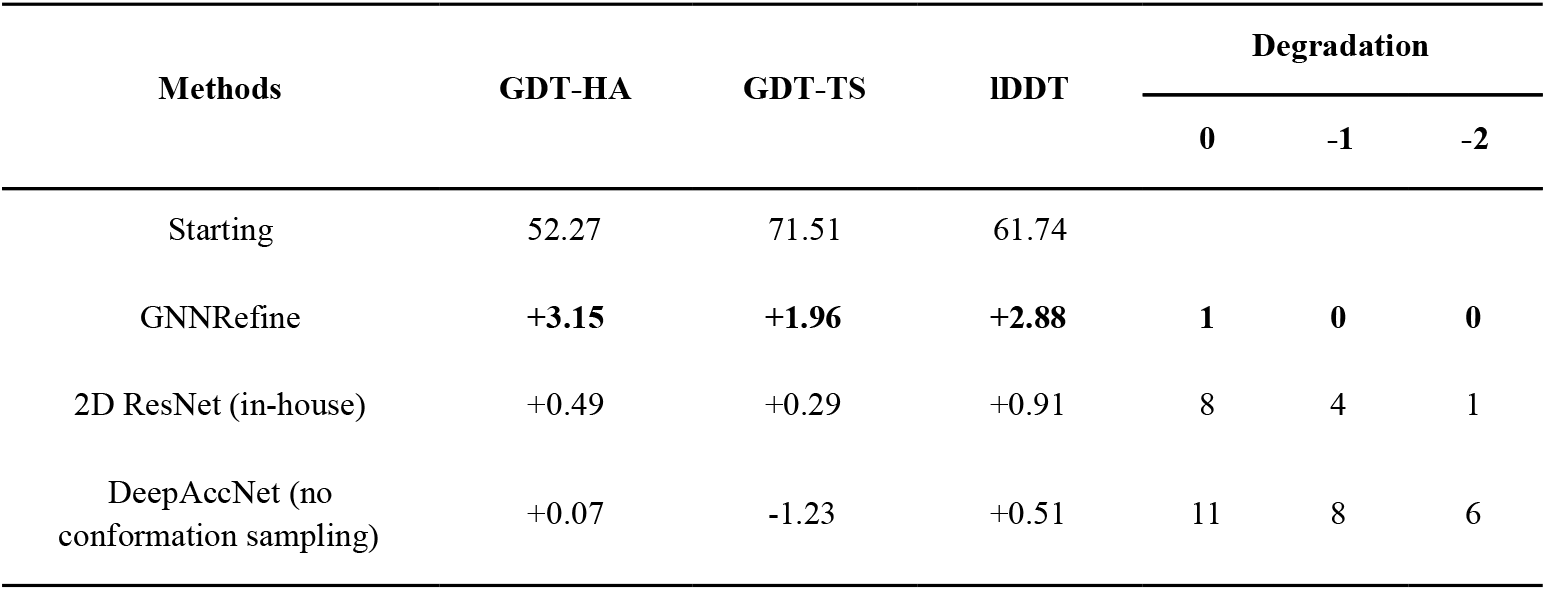
Performance of GNN-based and ResNet-based methods on the CASP13 refinement targets

| Methods | GDT-HA | GDT-TS | IDDT | Degradation |  |  |
| --- | --- | --- | --- | --- | --- | --- |
|  |  |  |  | 0 | -1 | -2 |
| Starting | 52.27 | 71.51 | 61.74 |  |  |  |
| GNNRefine | **+3.15** | **+1.96** | **+2.88** | **1** | **0** | **0** |
| 2D ResNet (in-house) | +0.49 | +0.29 | +0.91 | 8 | 4 | 1 |
| DeepAccNet (no conformation sampling) | +0.07 | -1.23 | +0.51 | 11 | 8 | 6 |

The underlying reason that GNN works better than ResNet for model refinement is that GNN is able to model the correlation of multiple residues more easily than ResNet. Most proteins have their radius of gyration proportional to the cube root of their length, so any two residues that are well separated along the primary sequence can be connected by a path in the protein graph shorter than the cube root of the protein length. As such, the correlation of multiple residues (spreading out in the distance matrix) can be modelled more effectively by (not so deep a) GNN, but not by a ResNet. That is, ResNet is good for inferring the initial inter-residue relationship and GNN is more suitable for refining it.

### Ablation study

To assess the contribution of individual factors to GNNRefine, we evaluate the GNNRefine models trained by different data and different features in Table 6. Supplementary Table S6 also shows the quality of predicted distance. In summary, the large training data, the inter-residue orientation and the DSSP-derived features are the three most important factors. The atom embedding on average does not provide useful information. Supplementary Table S7 shows the performance of iterative refinements by 5 GNNRefine models on the CASP13 targets, which demonstrates that the GNNRefine models trained on different datasets are complementary to each other. We have also predicted the CαCα distance, the NO distance, the inter-residue orientation and used them as restraints to build refined models, but did not observe significant improvement, as shown in Supplementary Table S8.

**Table 6.** Performance of GNN with different features and training data on the CASP13 refinement targets

| Features | Training data | GDT-HA | GDT-TS | IDDT | Degradation |  |  |
| --- | --- | --- | --- | --- | --- | --- | --- |
|  |  |  |  |  | 0 | -1 | -2 |
| All features | In-house | +3.15 | +1.96 | **+2.88** | **1** | **0** | **0** |
| All features | DeepAccNet data | +3.19 | +1.75 | +2.74 | 3 | 1 | 1 |
| All features | CASP models only | +1.42 | +0.92 | +1.35 | 8 | 6 | 3 |
| no Orientation | In-house | +2.21 | +1.28 | +2.26 | 4 | 2 | 0 |
| no Dihedral&SS&RSA | In-house | +2.53 | +1.67 | +2.31 | 2 | 0 | 0 |
| no AtomEmb | In-house | **+3.25** | **+2.03** | +2.57 | 2 | 0 | 0 |

## Discussion

This paper has presented a new method GNNRefine for protein model refinement. GNNRefine uses graph neural networks (GNN) to predict inter-residue distance distribution from an initial model and then feed the predicted distance information into PyRosetta FastRelax to build refined models. Since only limited conformation sampling is used, GNNRefine may refine models very quickly. Our study shows that even generating only 5 refined models from an initial model (within ∼15 minutes), GNNRefine can improve model quality almost as well as generating 50 refined models, and that GNNRefine can refine models as well as or better than some methods that use extensive conformation sampling. When conformation sampling is limited, GNNRefine works much better than ResNet for protein model refinement, because GNN may predict refined distance better than ResNet from an initial model. In our current implementation, GNNRefine does not make use of any sequence homologs. It will be interesting to investigate if GNNRefine may be further improved when multiple sequence alignment is used as its input. It will also be interesting to study if we can further improve GNNRefine by directly outputting atom coordinates from GNN instead of applying FastRelax to generate refined models, just like what AlphaFold2 has done.

## Methods

### Datasets

#### In-house training dataset

It includes the CASP7-12 models and the models built by RaptorX for the ∼29000 CATH domains. The CASP7-12 models are downloaded from http://predictioncenter.org/download_area/. There is only a small number (<600) of protein targets in CASP7-12. To increase the coverage, we select 28863 CATH domains (sequence identity <35%) released in March 2018^20^, and build on average 13 template-based and template-free models for each domain using our in-house protein structure prediction software RaptorX. In total, there are 29455 proteins with 509443 models in this training set. About 5% of the proteins and their decoys are randomly selected to form the validation set and the remaining decoys are used to form the training set. We generate 3 different training and validation splits and accordingly train three different GNNRefine models.

#### DeepAccNet training dataset^13^

It contains 7992 proteins (retrieved from the PISCES server^21^ and deposited to PDB by May 1, 2018) with 1104080 decoy models in total. Compared with our in-house training dataset, this dataset covers fewer protein targets (7992 v.s. 29455) but has many more decoy models for each target (∼138 v.s. ∼18). This set has a larger percentage of high-quality models. See Supplementary Figure S2 for the model quality distribution of these two datasets. We generate two different training and validation splits on this dataset and then train two different GNN models.

#### Test data

We use four test datasets to evaluate our method: the CASP13 refinement dataset, the CASP14 refinement dataset, the CAMEO dataset, and CASP13 FM dataset. The CASP13 refinement dataset includes 28 starting models in the CASP13 model refinement category^5^, excluding R0979 since it is an oligomeric target with three domains while our method is trained on individual domains. The CASP14 refinement dataset includes 37 starting models. The description of the CAMEO dataset and the CASP13 FM targets and their results are available at the Supplemental File. Note that all the training targets were released before May 1, 2018 and all the test targets were released after this date and thus, there is no overlap between our training and test datasets. The detailed information of our data is shown in the Supplementary Table S9.

### Feature extraction and graph definition

From a protein model, we derive two types of features: residue feature and residue pair feature. The residue feature includes sequential and structural properties of a residue: 1) one-hot encoding of the residue (i.e., a binary vector of 21 entries indicating its amino acid type); 2) the relative position of the residue in its sequence calculated as *i/L* (where *i* is the residue index and *L* is the sequence length); 3) dihedral angle (in radian), secondary structure (3-state), and relative solvent accessibility calculated by DSSP^22^; 4) one-hot encoding and relative coordinates of heavy atoms in the residue. The one-hot encoding is a four-dimensional vector representing four atom types (C, N, O and S) and the relative coordinate is a three-dimensional vector defined as: (*x* − *x*_α_, *y* − *y*_α_, *z* − *z*_α_), where (*x, y, z*) is a heavy atom’s coordinate and (*x*_α_, *y*_α_, *z*_α_) is the Cα atom’s coordinate.

The residue pair feature is derived for a pair of residues with Euclidean distance less than 10Å, including: 1) spatial distances of three atom pairs (CαCα, C_β_C_β_ and NO) scaled by 0.1; 2) three types of inter-residue orientation (ω, θ dihedrals and φ angle) defined in trRosetta^4^; 3) the sequential separation of the two residues (i.e. the absolute difference between the two residue indices), which is discretized into 9 bins ([1, 2, 3, 4, 5, 6-10, 11-15, 16-20, >20]) and represented by one-hot encoding. All these features are summarized in Supplementary Table S10.

We represent an initial protein model as a graph, in which one node represents a residue and one edge represents a chemical bond or a contact between two residues. We say there is a contact between two residues if their C_β_ Euclidean distance is no more than 10Å. It should be noted that this protein graph is equivariant (or symmetric) with respect to rotation and translation of atomic coordinates in the 3D space^23^.

### GNNRefine architecture and training

Our GNN model contains an atom embedding layer, 10 message passing layers, and an output layer. The dimensions of the atom embedding, edge feature and node feature are all 256. As shown in Figure 5A, the atom embedding layer is used to extract the local structure information for each residue. Its input is the one-hot encoding of an amino acid and the relative coordinates of heavy atoms in the residue, and its output is the atom embedding with a fixed dimension. The atom embedding is concatenated with other residue features to form the input feature of one residue (i.e. node feature in the graph). Each message passing layer consists of a message block for edges and a reduce block for nodes. The message block for edges updates edge features and obtains edge attention values (Figure 5B) and the reduce block for nodes updates node features (Figure 5C).

**Fig. 5.**
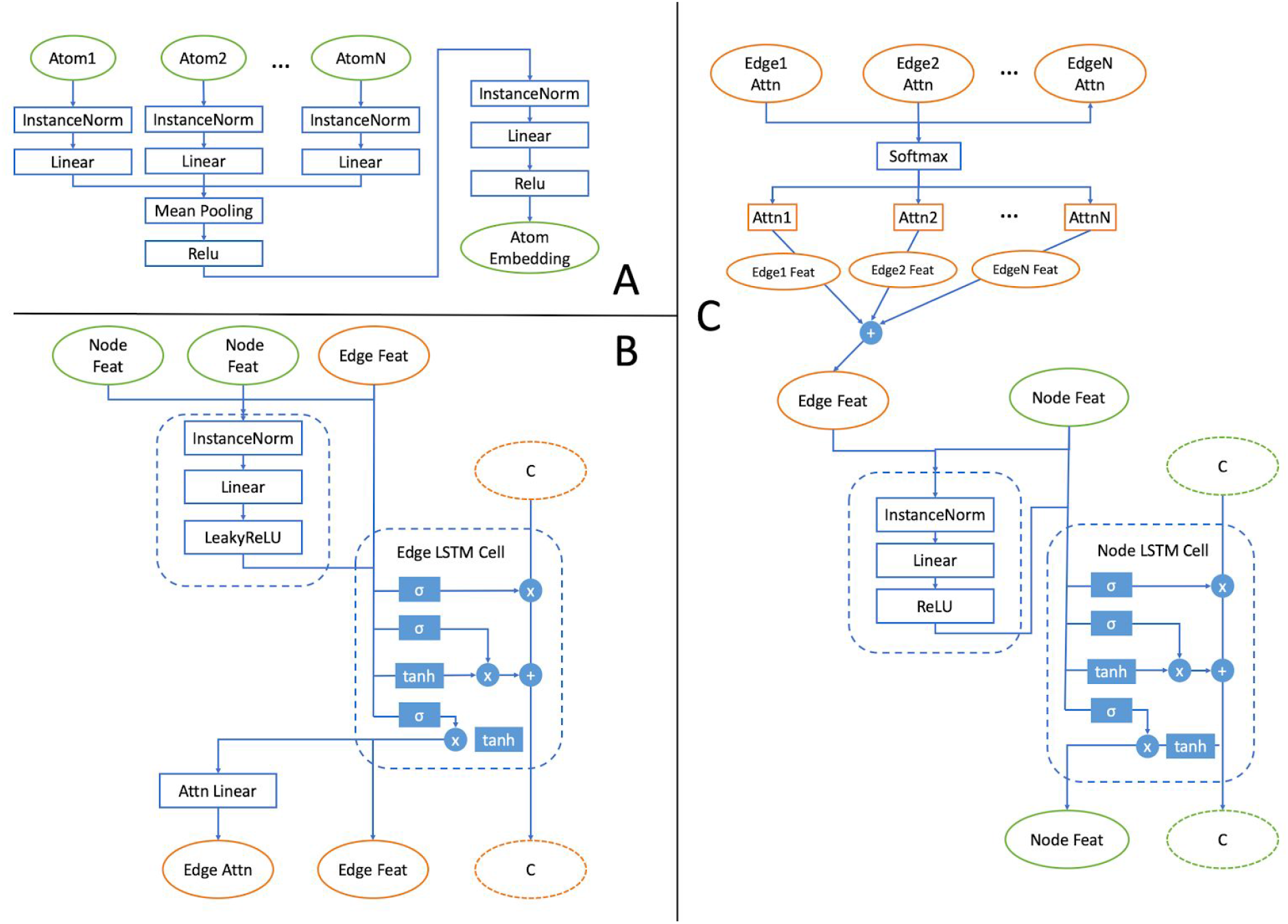
Detailed architectures of the atom embedding module and message passing blocks for edges and nodes. A. The atom embedding module; B. The message block for edges; C. The reduce block for nodes.

For each edge, the inputs of its message block are the features of the two nodes connected by the edge and the edge feature itself. All these features go through an instance norm layer, a linear layer, and a LeakyReLU layer to generate an intermediate edge feature, which then goes through an LSTM cell to obtain the new edge feature. For each LSTM cell, its input is the intermediate edge feature, its hidden state is the output of its preceding LSTM cell, and its cell state is updated from its preceding LSTM cell (the cell state of the first LSTM cell is initialized to 0). The LSTM cell may help to capture the long-term dependency across layers, which enables us to build a deeper GNN^24^. The new edge feature also goes through a linear attention layer to obtain the attention value of the edge. For each node, the inputs of its reduce block are the reduced edge feature and the node feature. The reduced edge feature is a linear combination of all edge features weighted by their respective attention values. Similar to an edge block, the features go through an instance norm layer, a linear layer, and a ReLU layer to generate an intermediate node feature, and then the intermediate node feature together with the initial node feature and its preceding LSTM cell state pass through an LSTM cell to obtain the new node feature and new cell state.

The output layer uses a linear layer and a softmax function to estimate the distance probability distribution based on the edge feature. The distance probability distribution is a 37-dimensional vector with 36 bins representing the distances from 2 to 20 Å (0.5Å each) and one bin indicating the distance >20Å, as presented in trRosetta^4^. To evaluate the refined model quality, we train a GNN-based quality assessment model which uses the node feature to predict the global lDDT and residue-wise lDDT simultaneously.

To fit the deep model to a GPU with limited memory, when a protein has more than 400 residues, a sub-structure of 400 consecutive residues is randomly sampled. We implement GNNRefine with DGL^25^ for PyTorch^26^ and train it using the Adam optimizer with parameters: β1=0.9 and β2=0.999. We set the initial learning rate to 0.0001 and divide it by 2 every 5 epochs. One minibatch has 16 protein models. We use the cross-entropy loss to train GNNRefine at most 15 epochs and select the model with the minimum loss on the validation data as our final model.

### Building refined models

Building a refined full-atom model by FastRelax^18^ consists of the following steps: 1) use the initial model to initialize the pose in PyRosetta; 2) convert the predicted distance probability distribution into distance potential using the DFIRE^27^ reference state and then add it onto the pose as restraints; 3) conduct full-atom relaxation, side-chain packaging and gradient-based energy minimization with the built-in *ref2015* ^28^ scoring function. The GNN model training and the FastRelax energy minimization may be trapped into local minima. To deal with this, we train 5 GNN models using five different training and validation data splits. Then we generate 10 refined models from an initial model using distance predicted by one GNN model and keep the lowest-energy one. We continue to refine this lowest-energy model using another GNN model until all 5 GNN models are applied. That is, in total we generate only 50 refined models and there are 5 lowest-energy refined models, which are then ranked by a GNN-based global model quality assessment (QA) method. This QA method is trained to predict global and local lDDT by the same set of data used to train GNNRefine. We also tested the strategy of generating only 1 refined model from an initial model by each of the 5 GNNRefine models sequentially. It turns out that this strategy has refinement accuracy almost as well as generating 10 refined models from an initial model by a single GNNRefine model.

## Acknowledgements

The authors are grateful to Prof. David Baker’s team including Hahnbeom Park who provided us the DeepAccNet training data. This work is supported by National Institutes of Health grant R01GM089753 to J.X. and National Science Foundation grant DBI1564955 to J.X. The funders had no role in study design, data collection and analysis, decision to publish, or preparation of the manuscript.

## Author contributions

X.J. conceived the research, developed the GNNRefine, and carried out the benchmarking experiments. J.X. built the in-house training data and guided the research. X.J. and J.X. analyzed the results and wrote the manuscript.

## Competing interests

The authors declare no competing interests.

## Notes

### Competing Interest Statement

The authors have declared no competing interest.

### Summary of Updates

1. corrected the description of the message block for edges. 2. added citations for the FastRelax and ref2015 scoring function.

