## Supplementary information for "Fast and effective protein model refinement by deep graph neural networks"

Xiaoyang Jing, Jinbo Xu

Toyota Technological Institute at Chicago, Chicago, IL 60637, USA

#### Performance on the CASP14 targets

Table S1. Performance on the CASP14 refinement targets when the AlphaFold2 models are excluded.

| **Methods** | **GDT-HA** | **GDT-TS** | **lDDT** | **Degradation** | | |
|---|---|---|---|---|---|---|
| | | | | **0** | **-1** | **-2** |
| Starting | 53.32 | 72.54 | 63.92 | | | |
| FEIG | +4.89 | +3.68 | +3.61 | 6 | 4 | 2 |
| BAKER | +2.67 | +0.84 | +2.28 | 8 | 8 | 7 |
| GNNRefine | 2.03 | +1.48 | +1.50 | **5** | 3 | **2** |
| GNNRefine[1] | +2.57 | +1.92 | +1.43 | 6 | **2** | **2** |
| FEIG-S | **+4.21** | **+2.93** | **+3.23** | **5** | 4 | 3 |
| Seok-server | +0.59 | +0.08 | +0.62 | 10 | 4 | **2** |
| Bhattacharya-Server | -0.14 | -0.16 | +0.48 | 15 | 9 | **2** |
| MUFOLD | -11.96 | -14.80 | -10.64 | 22 | 22 | 19 |

1. At each iteration, only one refined model is generated, i.e. in total only 5 refined models are generated by GNNRefine for each refinement target.

#### Performance on the CAMEO targets

Table S2. Performance of standalone software on the CAMEO targets

| Methods | GDT-HA | GDT-TS | lDDT | Degradation | | |
|---|---|---|---|---|---|---|
| | | | | **0** | **-1** | **-2** |
| Starting | 45.55 | 63.44 | 60.87 | | | |
| GNNRefine | **+1.99** | **+1.20** | **+2.23** | **39** | 21 | 12 |
| GalaxyRefine | +0.03 | +0.09 | +0.96 | 98 | 61 | 37 |
| ModRefiner-100 | -0.02 | -0.02 | +0.44 | 90 | **11** | **3** |
| ModRefiner-50 | -0.07 | -0.05 | +0.44 | 95 | 27 | 7 |
| ModRefiner-0 | -0.72 | -0.83 | +0.25 | 128 | 59 | 25 |
| FastRelax | -2.30 | -2.28 | -0.19 | 139 | 121 | 92 |

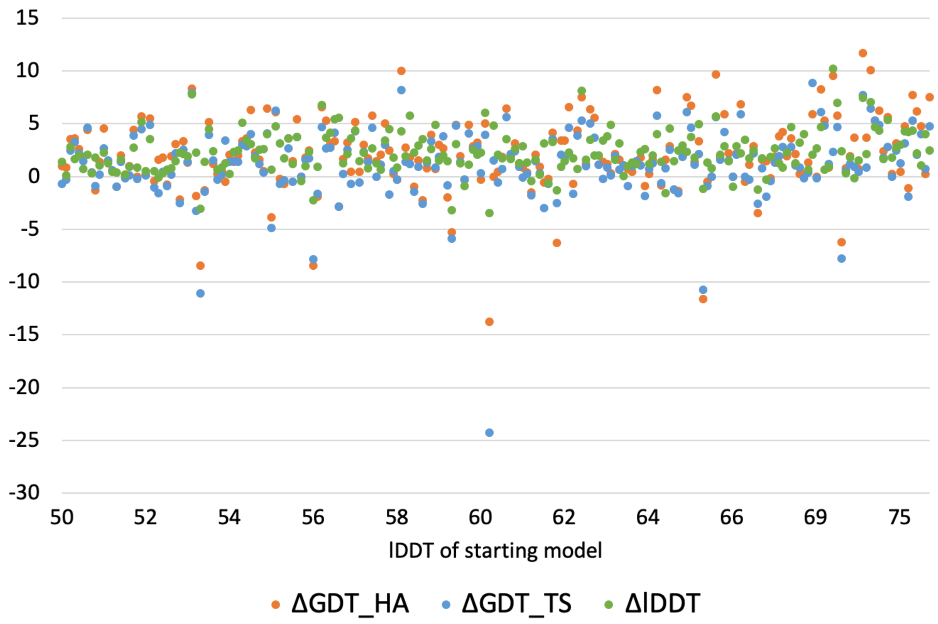


Fig. S1. The correlation between the starting model quality (of the CAMEO set) and the improvement by GNNRefine. The data are sorted from low to high according to the lDDT of their starting model.

Table S3. The quality of starting models vs. the quality improvements by GNNRefine on the CAMEO targets

| lDDT of starting model | Number of Models | Improvement | | |
|---|---|---|---|---|
| | | ΔGDT-HA | ΔGDT-TS | ΔlDDT |
| >50 | 208 | +1.99 | +1.20 | +2.23 |
| >60 | 100 | +2.39 | +1.51 | +2.38 |
| >70 | 21 | +4.09 | +2.60 | +3.15 |

| | | | | |
|---|---|---|---|---|
| >80 | 2 | +3.86 | +2.74 | +3.23 |

### Performance on the in-house CASP13 FM models

Table S4. Performance on our in-house decoys for the CASP 13 FM targets[1]

| Models | GDT-HA | | GDT-TS | | lDDT | | Degradation | | |
|---|---|---|---|---|---|---|---|---|---|
| | Starting | Refined | Starting | Refined | Starting | Refined | 0 | -1 | -2 |
| Max[2] | 40.46 | **+3.44** | 59.27 | **+2.79** | 52.02 | **+6.43** | 1 | 0 | 0 |
| Cluster10[3] | 32.38 | **+2.13** | 48.49 | **+2.01** | 46.93 | **+4.20** | 9 | 5 | 1 |
| Mean[4] | 34.85 | **+1.18** | 51.81 | **+0.71** | 48.24 | **+3.87** | 8 | 1 | 1 |

1. Only one refined model was generated for each decoy model.

2. The quality of the best model (in terms of GDT-HA) in the model pool for each target.

3. All models of a target are clustered into 10 clusters by SPICKER[1]. This row shows the mean quality of the 10 cluster centroids.

4. The mean quality of all models for each target.

### Distance prediction by GNN, in-house ResNet and DeepAccNet

Table S5. The improvement of predicted distances over the starting model by different methods on CASP13 targets

| Methods | Top L Contact Precision | | lDDT | |
|---|---|---|---|---|
| | Medium+Long | Long | Medium+Long | Long |
| GNN | **+6.99** | **+2.89** | **+3.26** | **+3.59** |
| 2D ResNet (in-house) | +4.83 | +1.06 | -1.07 | -1.16 |
| DeepAccNet DistPot[1] | -39.19 | -30.14 | -5.38 | -5.14 |

1. The predicted distance by DeepAccNet is derived from its distance potential. For each residue pair, the distance with the lowest potential is used as its predicted distance.

#### Ablation Study

Table S6. The improvement of predicted distance by different GNNRefine models on CASP13 targets

| Features | Training data | Top L Contact Precision | | lDDT | |
|---|---|---|---|---|---|
| | | Medium+Long | Long | Medium+Long | Long |
| All features | In-house | **+7.52** | +3.48 | +2.98 | +3.10 |
| All features | DeepAccNet data | +6.43 | +2.59 | +3.15 | +3.60 |
| All features | CASP models only | +3.94 | +1.09 | +0.55 | +1.11 |
| no Orientation | In-house | +7.38 | +2.54 | +2.15 | +2.22 |
| no Dihedral&SS&RSA | In-house | +6.99 | +2.89 | **+3.26** | **+3.59** |
| no AtomEmb | In-house | +7.40 | **+4.11** | +3.14 | +3.09 |

Table S7. Iterative refinement on the CASP13 targets using 5 different GNNRefine models trained on different datasets. Model 1, 2 and 3 are trained by 3 different splits of our in-house data and DAN 1 and 2 are trained by 2 different splits of the DeepAccNet data.

| Methods | GDT-HA | GDT-TS | lDDT | GDT-HA Degradation | | | lDDT Degradation |
|---|---|---|---|---|---|---|---|
| | | | | 0 | -1 | -2 | |
| Model 1 | +3.15 | +1.96 | +2.88 | 1 | 0 | 0 | 0 |
| + Model 2 | +3.40 | +2.16 | +3.09 | 3 | 1 | 0 | 0 |
| + Model 3 | +3.47 | +2.12 | +3.14 | 3 | 2 | 0 | 0 |
| + DAN 1 | +3.99 | +2.32 | +3.38 | 3 | 0 | 0 | 0 |
| + DAN 2 | +4.10 | +2.34 | +3.41 | 2 | 0 | 0 | 0 |

Table S8. Model refinement using restraints predicted for different atom types on the CASP13 targets

| Restraint | GDT-HA | GDT-TS | lDDT | GDT-HA Degradation | | |
|---|---|---|---|---|---|---|
| | | | | 0 | -1 | -2 |
| CbCb[1] | +3.15 | +1.96 | +2.88 | 1 | 0 | 0 |
| CbCb[2] | +2.94 | +1.73 | +2.55 | 3 | 3 | 0 |
| CaCa[2] | +3.09 | +1.95 | +2.69 | 3 | 2 | 1 |
| NO[2] | +1.17 | +0.47 | +1.06 | 7 | 2 | 2 |
| CaCa & CbCb[2] | +3.17 | +1.99 | +2.73 | 2 | 2 | 1 |

| | | | | | | |
|---|---|---|---|---|---|---|
| CaCa & CbCb & NO[2] | +2.52 | +1.45 | +2.02 | 4 | 3 | 0 |
| CbCb[3] | +2.76 | +1.56 | +2.45 | 4 | 3 | 1 |
| CbCb & Orientation[3] | +2.76 | +1.60 | +2.28 | 4 | 2 | 1 |

1. GNN model trained to predict CbCb distance only.
2. GNN model trained to predict CaCa, CbCb, and NO distances simultaneously.
3. GNN model trained to predict CbCb distance and orientation (ω, θ dihedrals and φ angle) simultaneously.

### Dataset description

Table S9. The number of protein targets and models of different datasets

| Dataset | Source | #Targets | | #Decoy or starting models | |
|---|---|---|---|---|---|
| In-house training data | CASP 7-12 | 592 | 29455 | 125045 | 509443 |
| | CATH | 28863 | | 384398 | |
| DeepAccNet training data | PISCES | 7992 | | 1104080 | |
| Test | CASP13 | 28 | | 28 | |
| | CASP14 | 37 | | 37 | |
| | CAMEO | 208 | | 208 | |
| | CASP13 FM | 28 | | 4193 | |

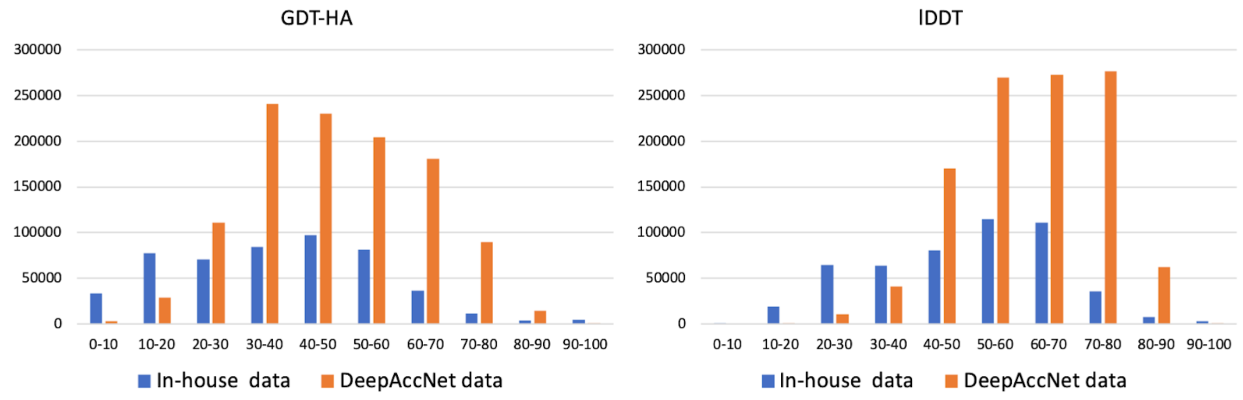


Fig. S2. The decoy quality distribution of the training data.

#### The CAMEO dataset and the CASP13 FM dataset

The CAMEO dataset includes 208 starting models from CAMEO[2]. To build this set, we download predicted protein models for all the CAMEO hard targets released between May 1st 2018 and May 1st 2020 from https://www.cameo3d.org/sp/. Then we keep only the targets with sequence lengths in [50, 500] and native structures containing at least 80% of sequence residues. Next, following the rule of CASP we select the best-predicted models (with the highest GDT-HA) for each target as starting models, and only keep the starting models with lDDT >50. For the CASP13 FM dataset, there are 28 targets corresponding to 32 FM domains defined by CASP13[3]. For each target we build ~150 decoys as starting models by our in-house template-free modeling method RaptorX-Contact[4].

#### Summary of Input Features

Table S10. Summary of input features

| Type | Feature | Dimension |
|---|---|---|
| Residue | One-hot encoding of residue | 21 |
| | rPosition | 1 |
| | Dihedral, SS3 and RSA calculated by DSSP | 6 |
| | One-hot encoding and relative coordinate of heavy atoms | 7 |
| Residue pair | Distance ($C\alpha C\alpha$, $C_\beta C_\beta$ and NO) | 3 |
| | Orientation ($\omega$, $\theta$ and $\varphi$) | 3 |
| | Sequential separation | 9 |
